# Temporal stability of sex ratio distorter prevalence in natural populations of the isopod *Armadillidium vulgare*

**DOI:** 10.1101/2023.11.27.568845

**Authors:** Sylvine Durand, Romain Pigeault, Isabelle Giraud, Anaïs Loisier, Nicolas Bech, Frédéric Grandjean, Thierry Rigaud, Jean Peccoud, Richard Cordaux

**Affiliations:** Laboratoire Ecologie et Biologie des Interactions, Equipe Ecologie Evolution Symbiose, Université de Poitiers, UMR CNRS 7267, Bât. B31, 3 rue Jacques Fort, TSA 51106, 86073 Poitiers Cedex 9, France; Laboratoire Biogéosciences, Université Bourgogne Franche-Comté, UMR CNRS 6282, 6 boulevard Gabriel, 21000 Dijon, France; Université Paris-Saclay, CNRS, IRD, UMR Évolution Génomes Comportement Écologie, 91190 Gif-sur-Yvette, France

**Keywords:** Sex ratio distorter, endosymbiont, *Wolbachia*, *f* element, sex determination, temporal dynamics

## Abstract

In the terrestrial isopod *Armadillidium vulgare*, many females produce progenies with female-biased sex ratios due to two feminizing sex ratio distorters (SRD): *Wolbachia* endosymbionts and a nuclear non-mendelian locus called the *f* element. To investigate the potential impact of these SRD on the evolution of host sex determination, we analyzed their temporal distribution in six *A. vulgare* populations sampled up to six times over 12 years, for a total of 29 time points. SRD distribution was heterogeneous among populations despite their close geographic locations, so that when one SRD was frequent in a population, the other SRD was rare. In contrast with spatial heterogeneity, our results overall did not reveal substantial temporal variability in SRD prevalence within populations, suggesting equilibria in SRD evolutionary dynamics may have been reached or nearly so. Temporal stability was also generally reflected in mitochondrial and nuclear variation. Nevertheless, in a population, a *Wolbachia* strain replacement coincided with changes in mitochondrial composition but no change in nuclear composition, thus constituting a typical example of mitochondrial sweep caused by endosymbiont rise in frequency. Rare incongruence between *Wolbachia* strains and mitochondrial haplotypes suggested the occurrence of intraspecific horizontal transmission, making it a biologically relevant parameter for *Wolbachia* evolutionary dynamics in *A. vulgare*. Overall, our results provide an empirical basis for future studies on SRD evolutionary dynamics in the context of multiple sex determination factors co-existing within a single species, to ultimately evaluate the impact of SRD on the evolution of host sex determination mechanisms and sex chromosomes.

## Introduction

Sex ratio distorters (SRD) are selfish genetic elements located on sex chromosomes or transmitted by a single sex, which skew the proportion of males and females in progenies towards the sex that enhances their own vertical transmission (Beukeboom and Perrin, 2014). They are found in a wide range of animal and plant species, of which they tremendously impact the ecology and evolution (Burt and Trivers, 2006; Werren, 2011). SRD are sources of genetic conflicts because they increase their own transmission at the expense of other genetic elements of the genome (Burt and Trivers, 2006; Werren, 2011; Beukeboom and Perrin, 2014). Genetic variants are therefore selected if they mitigate the fitness costs inflicted by SRD. Hence, female-biased sex ratios impose strong selective pressure, known as sex ratio selection, favouring genotypes producing more individuals of the under- represented sex (i.e. males) and ultimately restoring Fisherian (i.e. balanced) sex ratios. Thus, SRD may promote the evolution of host sex determination mechanisms (Burt and Trivers, 2006; Werren, 2011; Cordaux *et al*., 2011; Beukeboom and Perrin, 2014).

SRD are particularly well documented in arthropods, among which is the emblematic bacterial endosymbiont *Wolbachia* (Werren *et al*., 2008; Kaur *et al*., 2021). *Wolbachia* is a cytoplasmic, maternally inherited alpha-proteobacterium that often acts as a reproductive parasite by manipulating host reproduction in favor of infected females, thereby conferring itself a transmission advantage. In particular, *Wolbachia* has evolved the ability to induce female-biased sex ratios in host progenies through male killing, thelytokous (i.e. all female-producing) parthenogenesis and feminization (Werren *et al*., 2008; Cordaux *et al*., 2011; Hurst and Frost, 2015; Kaur *et al*., 2021).

Feminization, causing infected (and non-transmitting) genetic males to develop into (transmitting) phenotypic females, is mostly documented in terrestrial isopods (Martin *et al*., 1973). In the well-studied *Armadillidium vulgare*, chromosomal sex determination follows female heterogamety (ZZ males and ZW females) (Juchault and Legrand, 1972; Chebbi *et al*., 2019; Cordaux *et al*., 2021). In some *A. vulgare* populations, sex ratio is biased by *Wolbachia* bacteria or by a nuclear locus called the *f* element (Rigaud *et al*., 1997; Cordaux *et al*., 2011; Cordaux and Gilbert, 2017). Three *Wolbachia* strains are known to naturally occur in *A. vulgare*: *w*VulC, *w*VulM and *w*VulP (Rigaud *et al*., 1991; Cordaux *et al*., 2004; Verne *et al*., 2007). The *f* element results from the horizontal transfer of a large portion of a feminizing *Wolbachia* genome in the *A. vulgare* genome (Leclercq *et al*., 2016). The *f* element induces female development, as a W chromosome does, and it shows non-Mendelian inheritance, making it an SRD (Legrand and Juchault, 1984; Leclercq *et al*., 2016). In this species, sex ratio selection has resulted in the evolution of a masculinizing allele restoring balanced sex ratios, hence conferring resistance to feminization (Rigaud and Juchault, 1993). This nuclear locus represents a new male sex-determining locus, thereby effectively establishing a new male heterogametic system XY/XX. Thus, multiple sex determination factors co-exist in *A*. *vulgare* , ultimately caused by feminizing SRD (Juchault and Mocquard, 1993; Cordaux and Gilbert, 2017). Given the widespread distribution of *Wolbachia* infection in terrestrial isopods (Bouchon *et al*., 1998; Cordaux *et al*., 2012), it has been suggested that SRD may have contributed to the frequent turnovers of sex chromosomes recorded in this clade (Juchault and Mocquard, 1993; Juchault and Rigaud, 1995; Becking *et al*., 2017, 2019; Russell *et al*., 2021).

To elucidate the potential impact of SRD on the evolution of sex determination in terrestrial isopods, it is essential to clarify the intraspecific evolutionary dynamics of SRD such as *Wolbachia* and the *f* element. Previous studies have shown that: (i) *Wolbachia* and the *f* element are present at variable frequencies in *A. vulgare* populations, (ii) the *f* element is overall more frequent than *Wolbachia*, (iii) the two SRD usually do not co-occur at high frequency in populations, and (iv) mitochondrial haplotypes (mitotypes) are tightly linked to *Wolbachia* strains (suggesting stable maternal transmission), but not to the *f* element (Juchault *et al*., 1993; Durand *et al*., 2023). These results have provided insights into the spatial distribution of *Wolbachia* and the *f* element in *A. vulgare*; in the present study, we investigate their temporal dynamics.

Prior studies on endosymbiont temporal dynamics have shown that host population invasions can occur within just a few years, e.g. *Wolbachia* in *Drosophila simulans* (Turelli and Hoffmann, 1991) and *Eurema hecabe* (Miyata *et al*., 2024), and *Rickettsia* in *Bemisia tabaci* (Himler *et al*., 2011). Likewise, nuclear suppressors of SRD can rise in frequency very quickly, e.g. *Wolbachia* suppressor in *Hypolimna bolina* (Charlat *et al*., 2007). Similar patterns of rapid spread and evolution of resistance have been reported for a nuclear SRD, Paris Sex Ratio, in *D. simulans* (Helleu *et al*., 2019).

Endosymbiont temporal dynamics has seldom been studied on a longer time scale. Comparison of historical (i.e. museum) and contemporary sympatric populations of *H. bolina* highlighted fluctuations of *Wolbachia* frequency over periods of 73-123 years (Hornett *et al*., 2009). However, at an intermediate time scale (10-20 years), *Wolbachia* was found to decrease in frequency in *Acraea encedon* (Hassan *et al*., 2013), while it was stably maintained in *Eurema mandarina* (Kageyama *et al*., 2020) and *D. simulans* (Weeks *et al*., 2007; Carrington *et al*., 2011). In the case of *A. vulgare*, SRD temporal dynamics has previously been tackled by a single study (Juchault *et al*., 1992), in which *Wolbachia* prevalence was found to decrease concomitantly to an increase of *f* element prevalence in an *A. vulgare* population from Niort (western France) sampled at three time points over a period of 23 years (1963, 1973 and 1986). However, a single population was included in the study, thus limiting the breadth of its conclusions. Here we report an analysis of *Wolbachia* and *f* element distribution in six *A. vulgare* populations sampled up to six times over a period of 12 years, representing a total of 889 individuals from 29 time points. The studied populations were sampled in a narrow geographic area in western France, within 70 km of the Niort population (Juchault *et al*., 1992), to control for spatial dynamics. In contrast to most previous studies, here we analyzed SRD in the context of both mitochondrial and nuclear variation. Our results highlight an overall temporal stability of SRD distribution in *A. vulgare*, with few exceptions.

## Materials and Methods

A total of 889 *A. vulgare* individuals were analyzed, from six natural populations sampled in western France between 2003 and 2017 (Figure 1). DNA samples from Beauvoir-2017, Chizé-2017, Coulombiers-2017, Gript-2017, La Crèche-2017 and Poitiers-2015 were available from (Durand *et al*., 2023). All other individuals were collected by hand. Sex ratios in sampled individuals should not be considered as reflecting those of populations, due to possible selection biases by samplers.

**Figure 1.**
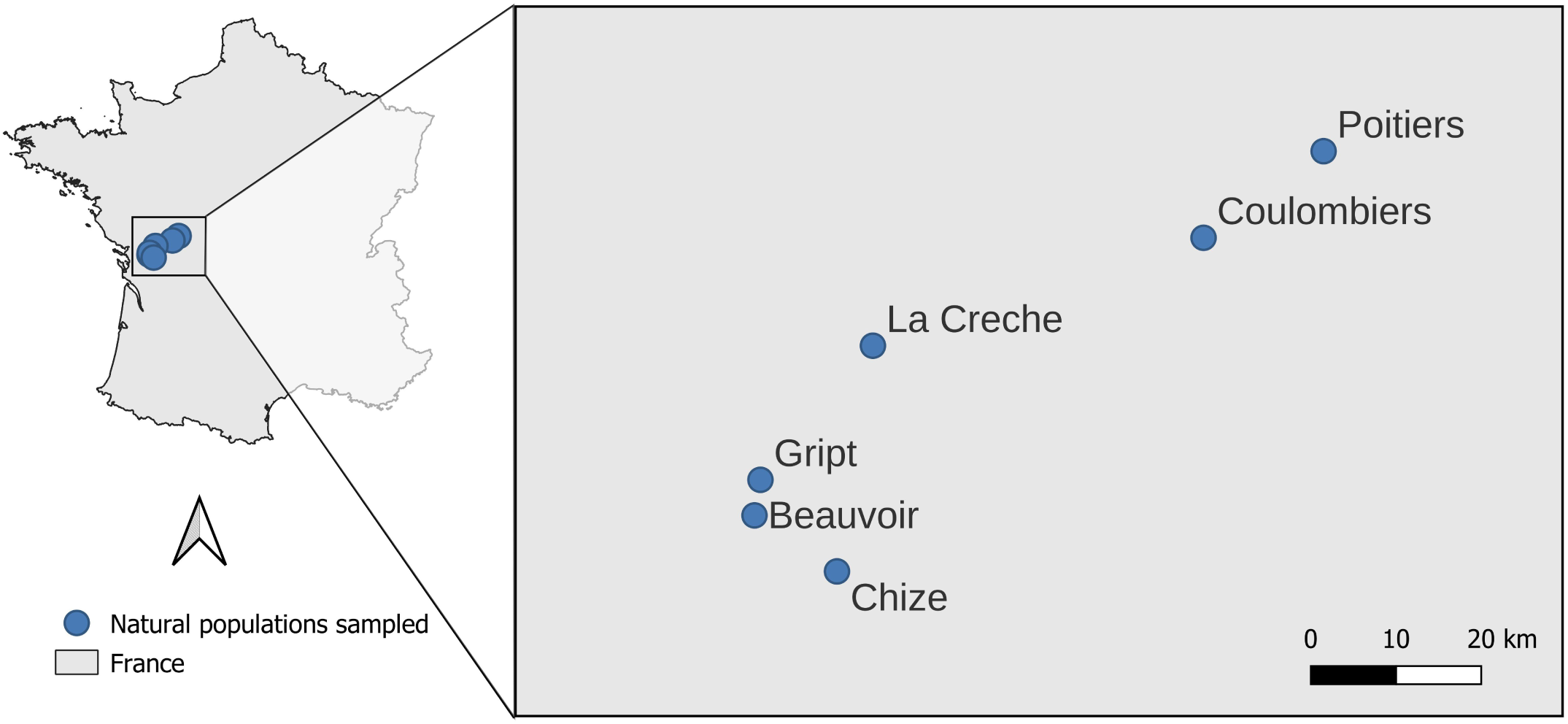
Map of France (left) showing the location of the six populations analyzed in this study and zoom in the relevant geographic area (right), indicating the relative positions and distances in kilometers (km) between populations.

Individuals were sexed and stored in alcohol or at -20°C prior to DNA extraction. Total genomic DNA of samples collected between 2003 and 2013 was extracted from gonads using phenol and chloroform (Kocher *et al*., 1989) and DNA of samples collected between 2014 and 2016 was extracted from the head and legs, as described previously (Leclercq *et al*., 2016).

Four molecular markers were used to assess the presence of *Wolbachia* and the *f* element in DNA extracts: *Jtel* (Leclercq *et al*., 2016), *wsp* (Braig *et al*., 1998), *recR* (Badawi *et al*., 2014) and *ftsZ* (Werren *et al*., 1995) (Table S1). While *Jtel* is specific to the *f* element, *wsp* and *recR* are specific to *Wolbachia*, and *ftsZ* is present in both the *f* element and *Wolbachia* (Leclercq *et al*., 2016). The presence or absence of these markers was assessed by PCR assays, as described previously (Leclercq *et al*., 2016). Different amplification patterns were expected for individuals with *Wolbachia* only (*Jtel- , wsp+*, *recR+*, *ftsZ+*), the *f* element only (*Jtel+, wsp-, recR-*, *ftsZ+*), both *Wolbachia* and the *f* element present (*Jtel+, wsp+, recR+*, *ftsZ+*) or both *Wolbachia* and the *f* element lacking (*Jtel-, wsp-, recR-*, *ftsZ-*). The few individuals exhibiting other amplification patterns were classified as “undetermined status”.

To characterize *Wolbachia* strain diversity, *wsp* PCR products were purified and Sanger sequenced using both forward and reverse primers by GenoScreen (Lille, France). Sequences from forward and reverse reads were assembled using Geneious v.7.1.9 to obtain one consensus sequence per individual.

To evaluate mitochondrial diversity, a ∼700 bp-long portion of the Cytochrome Oxidase I (*COI*) gene was amplified by PCR in all individuals (Folmer *et al*., 1994) (Table S1). PCR products were purified and Sanger sequenced as described above. Mitotype networks were built using the *pegas* package for R (Paradis, 2010).

To evaluate nuclear diversity, all individuals were genotyped at 22 microsatellite markers (Verne *et al*., 2006; Giraud *et al*., 2013) distributed in five multiplexes (Multiplex 1: Av1, Av2, Av4, Av5, Av9; Multiplex 2: Av6, Av3, Av8; Multiplex 3: AV0023, AV0056, AV0085, AV0086, AV0096; Multiplex 4: AV0002, AV0016, AV0018, AV0032, AV0099; Multiplex 5: AV0061, AV0063, AV0089, AV0128) (Table S2). PCR was performed using fluorescence-marked forward primers, as described previously (Durand *et al*., 2017). PCR fragments were separated by electrophoresis on an ABI 3730XL automated sequencer by Genoscreen (Lille, France). Alleles were scored using the software GeneMapper 3.7 (Applied Biosystems), each genotype being independently read by two people.

Of the 22 amplified microsatellite markers, Av4 and Av5 could not be scored because of multiple peaks, AV0096 and AV0128 did not amplify consistently, and AV0023 and AV0061 were monomorphic across the dataset. The *Genepop* package for R (Rousset, 2008) detected no linkage disequilibrium between the 16 remaining loci. The presence of null alleles was tested by using a combination of software, as recommended previously (Dąbrowski *et al*., 2014): Micro-Checker (Van Oosterhout *et al*., 2004), Cervus (Kalinowski *et al*., 2007) and ML-NullFreq (Kalinowski and Taper, 2006). As a result, AV0099 was discarded because it consistently presented hints of null alleles in many populations and sampling years. Individuals whose genotypes were available for fewer than 13 out of the 15 remaining markers were removed from the following analyses.

Hardy-Weinberg equilibrium was tested for each locus, locality and sampling year with an exact test using Markov chain with the *Genepop* package for R. The Fstat software v. 2.9.3.2 (Goudet, 2001) was used to calculate allelic richness and heterozygosity (based on a minimum of 3 individuals).

Genetic differentiation was estimated by computing Fst values for all pairs of populations (Weir and Cockerham, 1984) with Fstat. Significance was calculated using global tests implemented in Fstat with a level of significance adjusted for multiple tests using the standard Bonferroni correction. The longitudes and latitudes of the populations were used to calculate Euclidean distances between populations and to test for isolation by distance by correlating these geographical distances with the genetic distances. Significance of the correlation was tested at individual scale by using a Mantel test (with 9999 permutations) implemented in GENALEX v 6.2 (Peakall and Smouse, 2006). Genetic clusters were also delineated without a priori with a Bayesian, individual-based approach implemented in the software Structure (Pritchard *et al*., 2000). The admixture model was selected, as well as the option of correlated allele frequencies. The number of clusters (K) varied from 2 to 9. For each value of K, 20 independent runs were carried out, as recommended in (Evanno *et al*., 2005), with a total number of 100,000 iterations and a burn-in of 10,000. To determine the most likely value of K, the method described in (Evanno *et al*., 2005) was applied as implemented in Structure Harvester version 0.6.9 (Earl and vonHoldt, 2012). In addition, a Discriminant Analysis of Principal Components (DAPC) (Jombart *et al*., 2010) was performed on populations according to their sampling locality and year with the *adegenet* package (Jombart, 2008), to search for potential discrepancies across time points within populations.

Statistical analyses were carried out using the R software (v.4.2.1). The influence of sex, and population if appropriate, on SRD prevalence was analyzed using general linear models with a binomial error distribution. Maximal models, including all higher order interactions, were simplified by sequentially eliminating non-significant terms and interactions to establish a minimal model (Crawley, 2012). The significance of the explanatory variables was established using a likelihood ratio test, which is approximately distributed as a Chi-square distribution (Bolker, 2008). The significant Chi-squared values given in the text are for the minimal model, whereas non-significant values correspond to those obtained before the deletion of the variable from the model. Chi-square tests were used to compare the frequency of individuals infected by *Wolbachia* and the frequency of individuals carrying the *f*-element in the different populations. Figures were realized with *ggplot2* (Wickham *et al*., 2020). Results from Structure were processed with the program Distruct (Rosenberg, 2003) for graphical representation.

## Results

We tested the presence of *Wolbachia* and the *f* element in 889 individuals (627 females and 262 males) from six populations sampled at various time points between 2003 and 2017, representing a total of 29 sampling points (Tables 1 and S3). Among the 889 analyzed individuals, we failed to determine the status of 29 individuals. Of the remaining 860 individuals, 29.9% carried only the *f* element, 15.2% carried only *Wolbachia*, 0.6% carried both SRD and 54.3% carried none. Although sometimes present in males, both SRD were significantly more frequent in females than in males (*Wolbachia*: χ² = 73.59, p < 0.0001; *f* element: χ² = 41.03, p < 0.0001, Tables 1 and S3). *Wolbachia*- infected individuals carried one of the three previously known *Wolbachia* strains of *A. vulgare*: *w*VulC (n=22), *w*VulM (n=25) or *w*VulP (n=83).

Overall, both SRD were found at at least one time point in all six populations (Figure 2). However, the populations displayed substantial variation in the distribution of the two SRD. In Beauvoir, Chizé, Coulombiers and La Crèche populations, the *f* element was significantly more predominant (29-86% frequency) than *Wolbachia* (2-10%) (χ² > 36.91 and p < 10^-9^ in all four populations). Conversely, *Wolbachia* was significantly more frequent (63%) than the *f* element (2%) in Poitiers (χ² > 135.32, p < 10^-16^). Finally, both SRD were very rare in Gript (2-3%) and did not differ in frequencies (χ² = 0.10, p = 0.748). Remarkably, *f* element prevalence did not differ significantly across time points (spanning up to 12 years) in all four populations in which it was predominant (χ² = 0.002, p =0.968), although *f* element prevalence was significantly different among populations (χ² = 62.55, p < 0.0001). In these populations, *f* element prevalence was significantly higher in females than in males (χ² = 61.87, p < 0.0001) (Figure 2). In Poitiers, where *Wolbachia* remained globally frequent over time, females were also significantly more infected than males (χ² = 106.06, p < 0.0001). Interestingly, the frequency of the *w*VulC strain significantly decreased over time (χ² = 12.18, p < 0.0001) while the *w*VulP strain exhibited the opposite trend, and significantly so (χ² = 6.03, p = 0.014).

**Figure 2.**
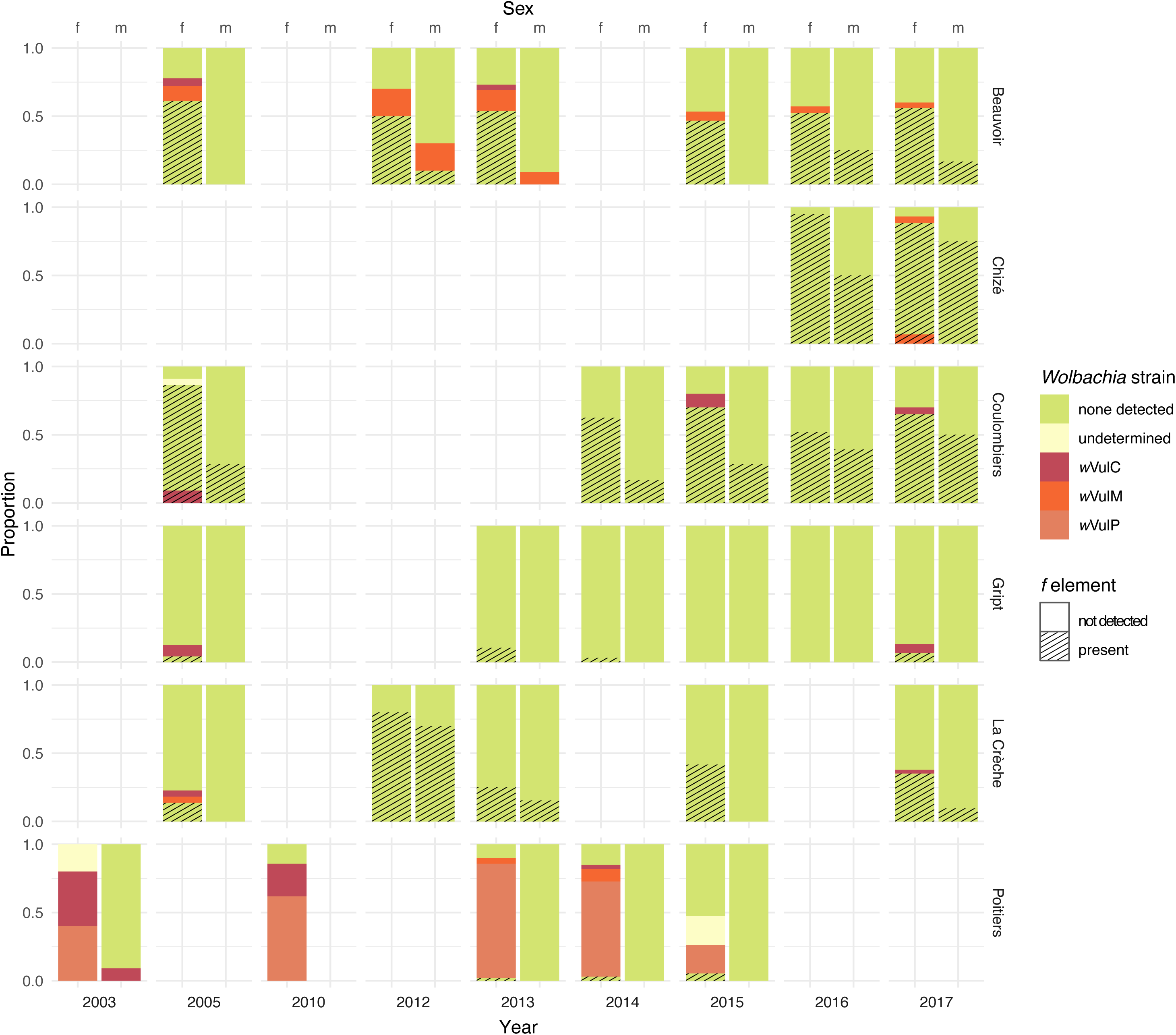
Prevalence of sex ratio distorters in *Armadillidium vulgare* males (m) and females (f) from six populations sampled at different time points.

Sequencing of the *COI* mitochondrial gene of 884 individuals identified 12 mitotypes, nine of which have previously been detected in *A. vulgare* populations (named I to VIII, and XII) (Durand *et al*., 2023) and three are newly described mitotypes (XXIV to XXVI; GenBank accession numbers OR074129 to OR074131, respectively) (Figure 3 and Table S3). There was an excellent congruence between *Wolbachia* strains and mitotypes, as previously reported (Verne *et al*., 2012; Durand *et al*., 2023). Indeed, individuals carrying *w*VulC were associated with either mitotype V or its close relatives (XII and XXVI), those carrying *w*VulP were associated with mitotype VII and those carrying *w*VulM were associated with mitotype II. Exceptions included two individuals infected by *w*VulM, which were associated with the distantly related mitotypes I and III. By contrast, the *f* element was found in eight different mitochondrial backgrounds (I to VI, VIII and XXV) distributed across the mitochondrial network.

**Figure 3.**
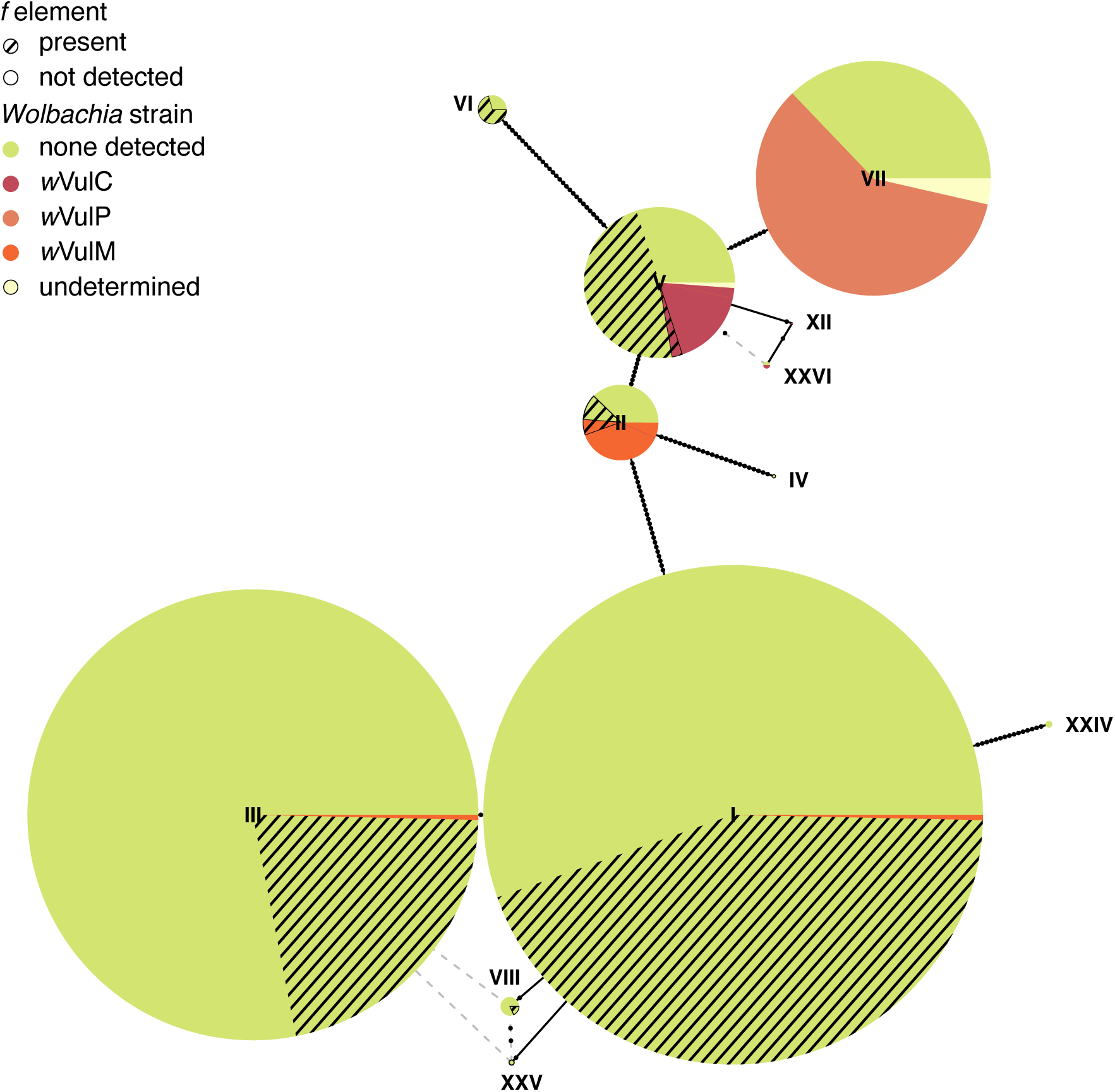
Mitotype network of 12 mitochondrial variants from six *Armadillidium vulgare* populations sampled at different time points. Each circle represents one mitotype and circle diameter is proportional to the number of individuals carrying the mitotype. Branch lengths connecting circles are proportional to divergence between mitotypes.

All time points considered, there was between three (in Chizé) and six (in Poitiers) mitotypes per population (Figures 3 and S1). Mitotype frequencies were globally stable across time points within populations (e.g. Coulombiers, Figure 4A), with the notable exceptions of La Crèche and Poitiers populations. In La Crèche, a shift in major mitotypes occurred between 2005 and 2012, with the increasing rarity of mitotype III and V being concomitant with the rise in frequency of mitotypes I and VIII, followed by stability since 2012 (Figure 4B). In Poitiers, the rise in frequency of mitotype VII (associated with *Wolbachia* strain *w*VulP) coincided with a relative decrease of *w*VulC-associated mitotypes V, XII and XXVI (Figure 4C).

**Figure 4.**
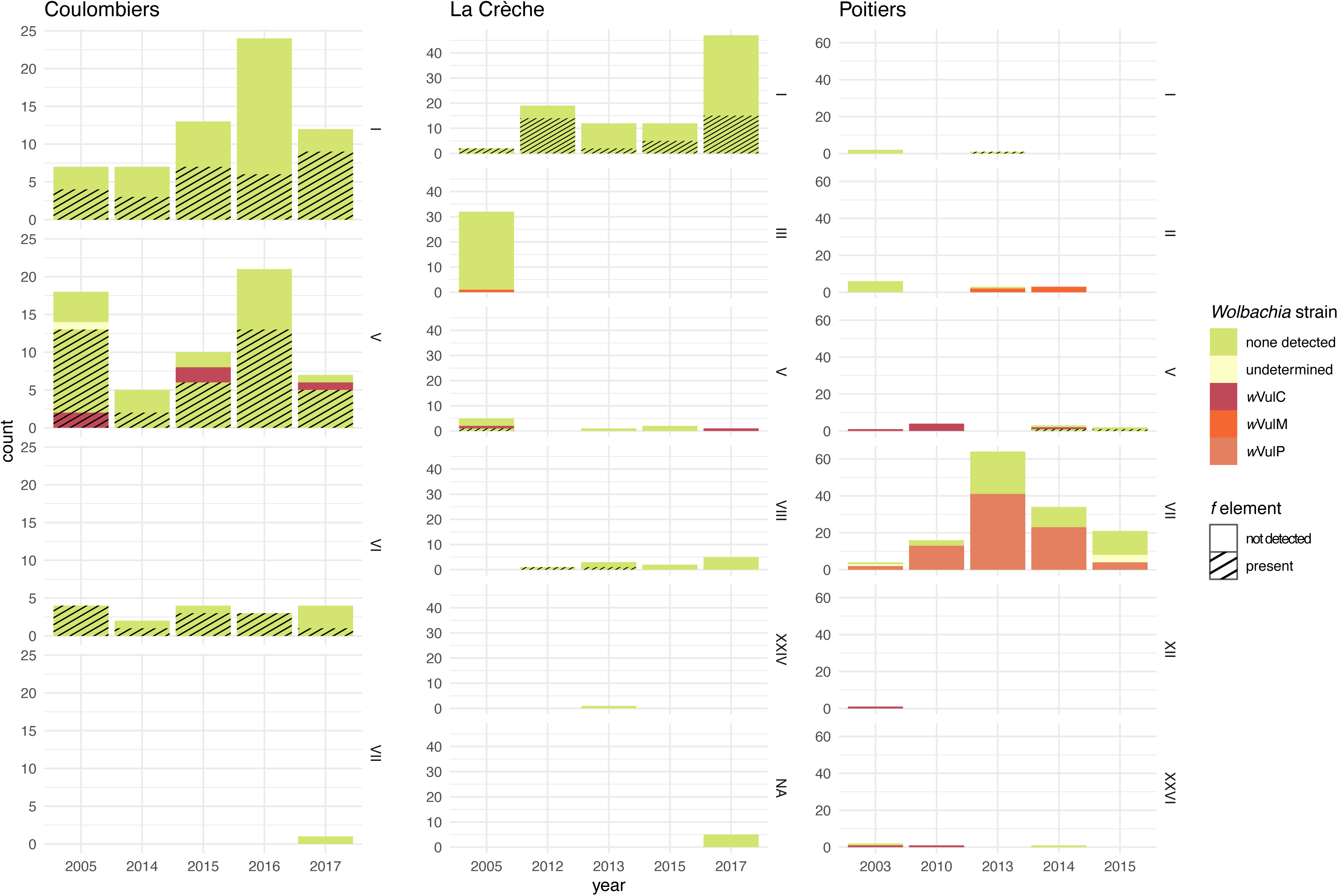
Variation in mitotype counts across years for three *Armadillidium vulgare* populations from Coulombiers, La Crèche and Poitiers. Prevalence of sex ratio distorters is coded as indicated in the key on the right-hand side of the figure.

To test if the patterns of SRD and mitochondrial variation were also reflected in the nuclear genome, we examined variation in 667 individuals with genotype information available for at least 13 out of the 15 retained microsatellite markers (see Materials and Methods), representing a total of 28 sampling points (Poitiers-2015 was discarded due to low genotyping success) (Table S3). None of the loci significantly departed from Hardy-Weinberg equilibrium for all sampling points. Allelic richness ranged from 1 to 4.78 and heterozygosity ranged from 0 to 0.91 (Table S4). Pairwise Fst values ranged from 0 to 0.091 (mean Fst = 0.035) (Table S5). The majority of Fst values between populations (200/321) were significant, suggesting the occurrence of genetic differentiation among populations. Consistently, a Mantel test evaluating the correlation between genetic and geographic distances revealed a significant isolation by distance (r^2^ = 0.034, p < 0.001). Such a signal of genetic structure was confirmed by Bayesian clustering analyses. Indeed, a delta-K analysis (Evanno et al., 2005) inferred that the best fit to the data was obtained for K=2 genetic clusters (Figure S2). A first genetic cluster mainly comprised individuals sampled in Poitiers and Coulombiers, and a second genetic cluster comprised those from Beauvoir, Chizé, La Crèche and Gript (Figure 5). In agreement with isolation by distance, this clustering pattern reflected the geographic distribution of populations, as the former cluster encompassed eastern-most populations and the latter cluster encompassed western-most populations (Figure 1). It is noteworthy that the low delta-K values obtained suggested poor convergence between the 20 independent runs, which may be the result of a previously highlighted effect of isolation by distance (Meirmans, 2012). By contrast, the clustering analysis did not highlight any obvious change in genetic structure between time points within populations. The DAPC also suggested a major population structuration (Figure 6), as the first component separated Poitiers and Coulombiers (locating towards the right side of the axis) from the other populations (locating towards the left side of the axis). It also supported overall homogeneity of populations across time points, apart from La Crèche-2005, which was separated from the other La Crèche time points in the second component of the DAPC. The exception of La Crèche-2005 was also reflected in the significant pairwise Fst values between time points within population, which were generally non-significant for the other populations (Table S5).

**Figure 5.**
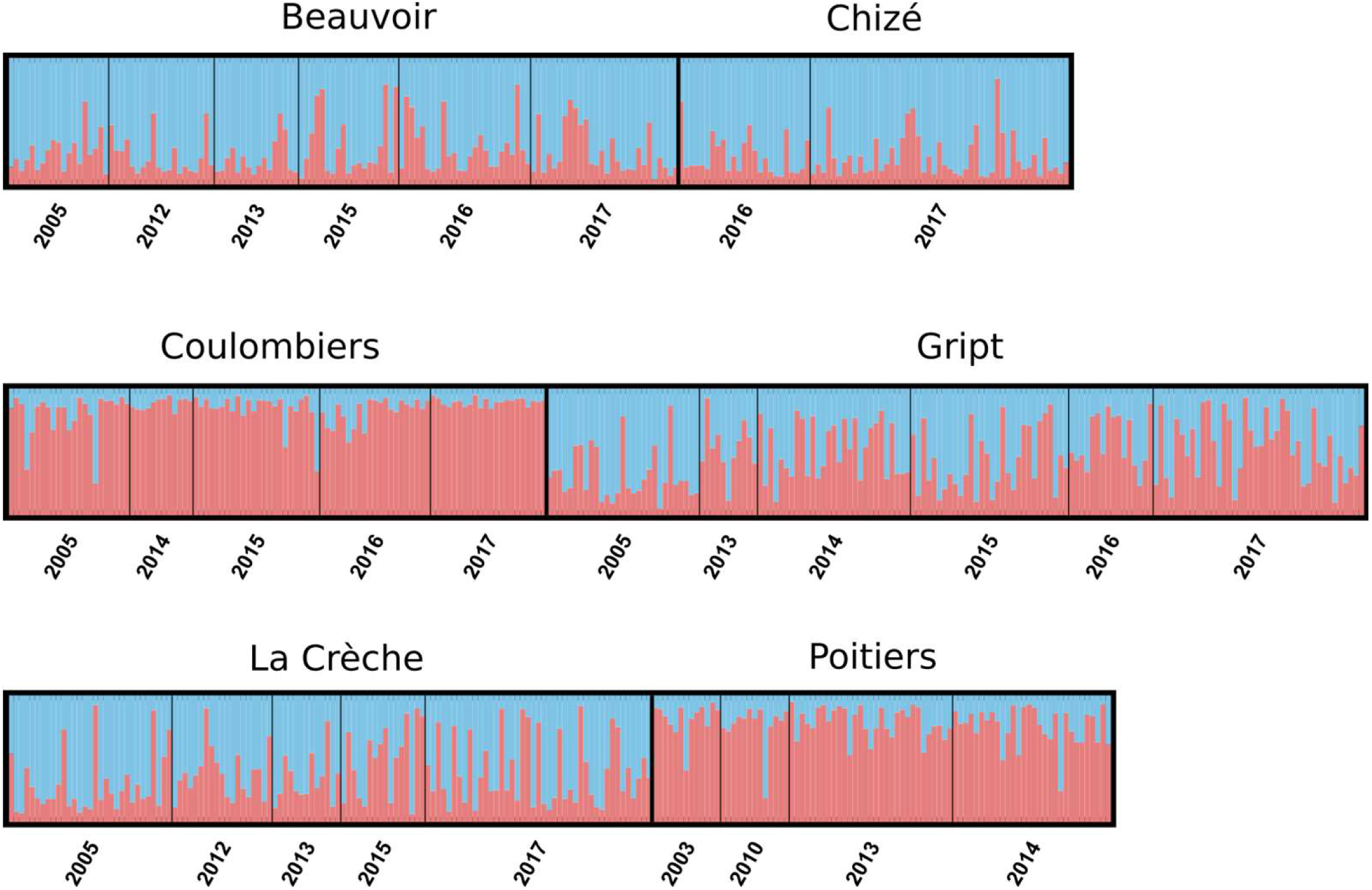
Assignment of individuals from six *Armadillidium vulgare* populations sampled at different time points to one of two genetic clusters (blue and pink colors) following Bayesian analysis. Each bar represents an individual, and the proportion of each color represents the probability of assignment to the corresponding cluster.

**Figure 6.**
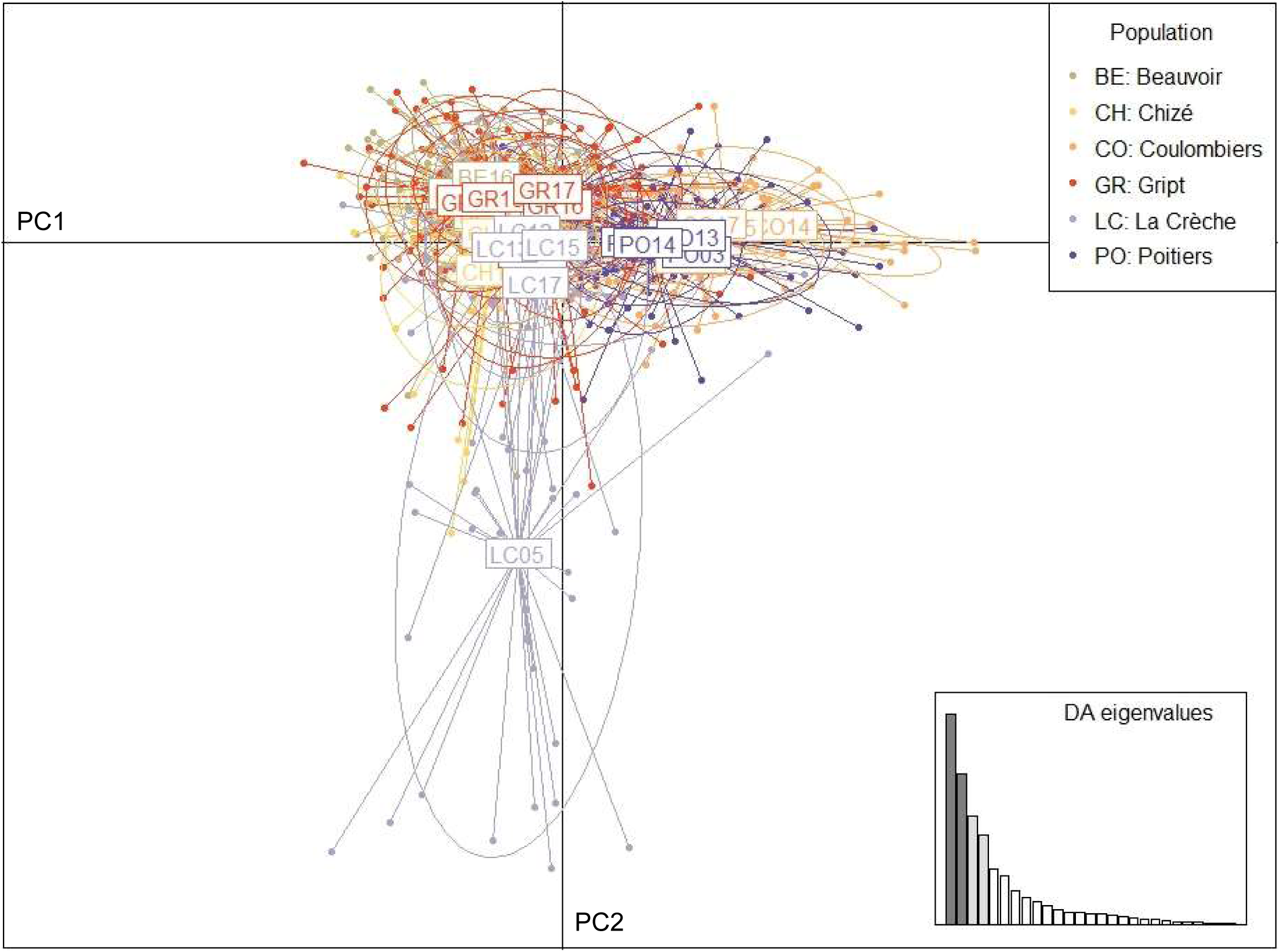
Discriminant Analysis of Principal Components scatterplot. The two axes represent the first two principal components (PC). Dots represent individuals. Each of the 28 sampling points presents a 95% inertia ellipse and is labeled with two letters indicating the population and the two last digits of the sampling year. The eigenvalues of the analysis (inset) show the relative amount of genetic structure captured by the principal components (the first two components are highlighted in dark grey).

**Table 1.**
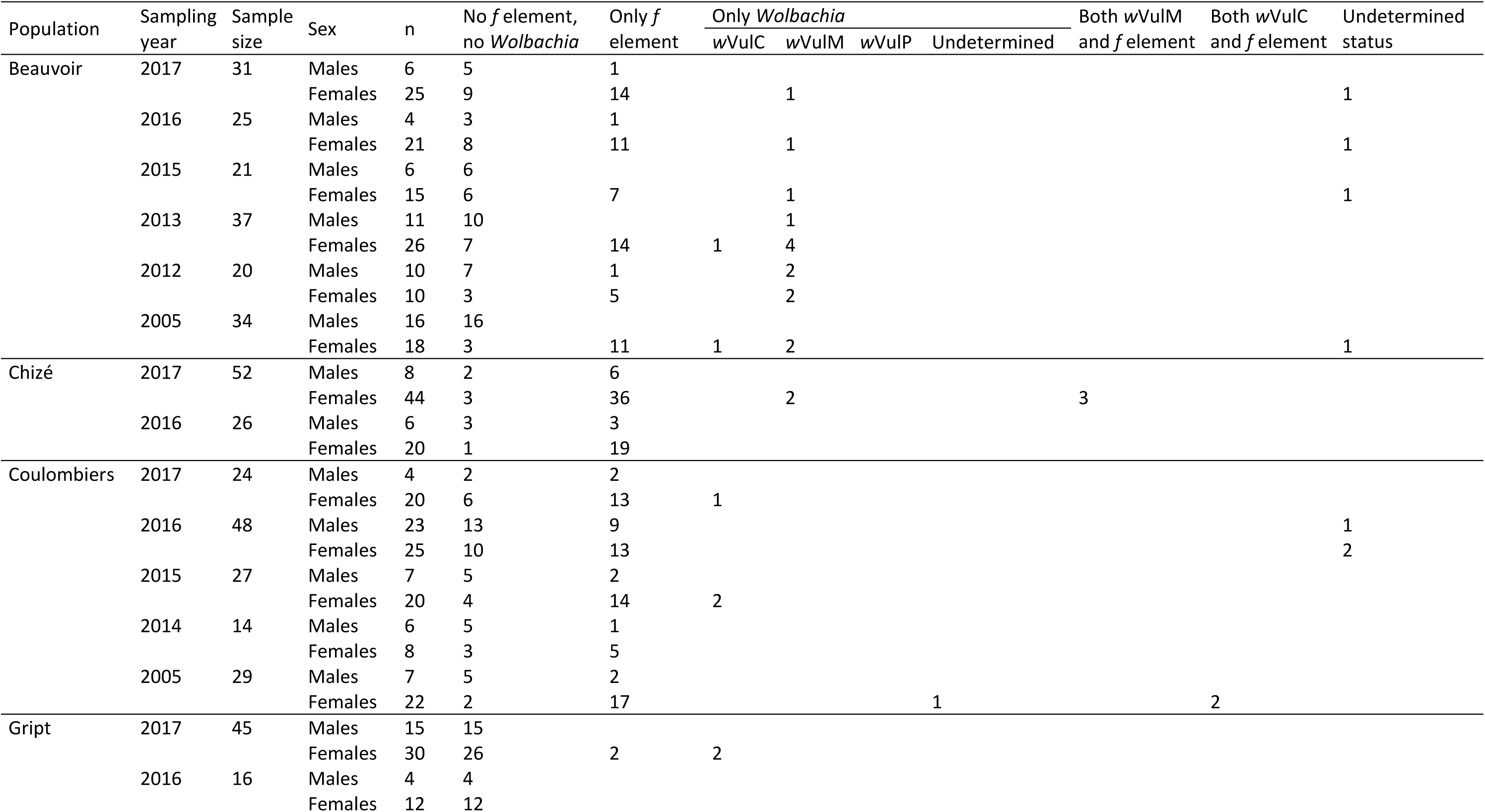
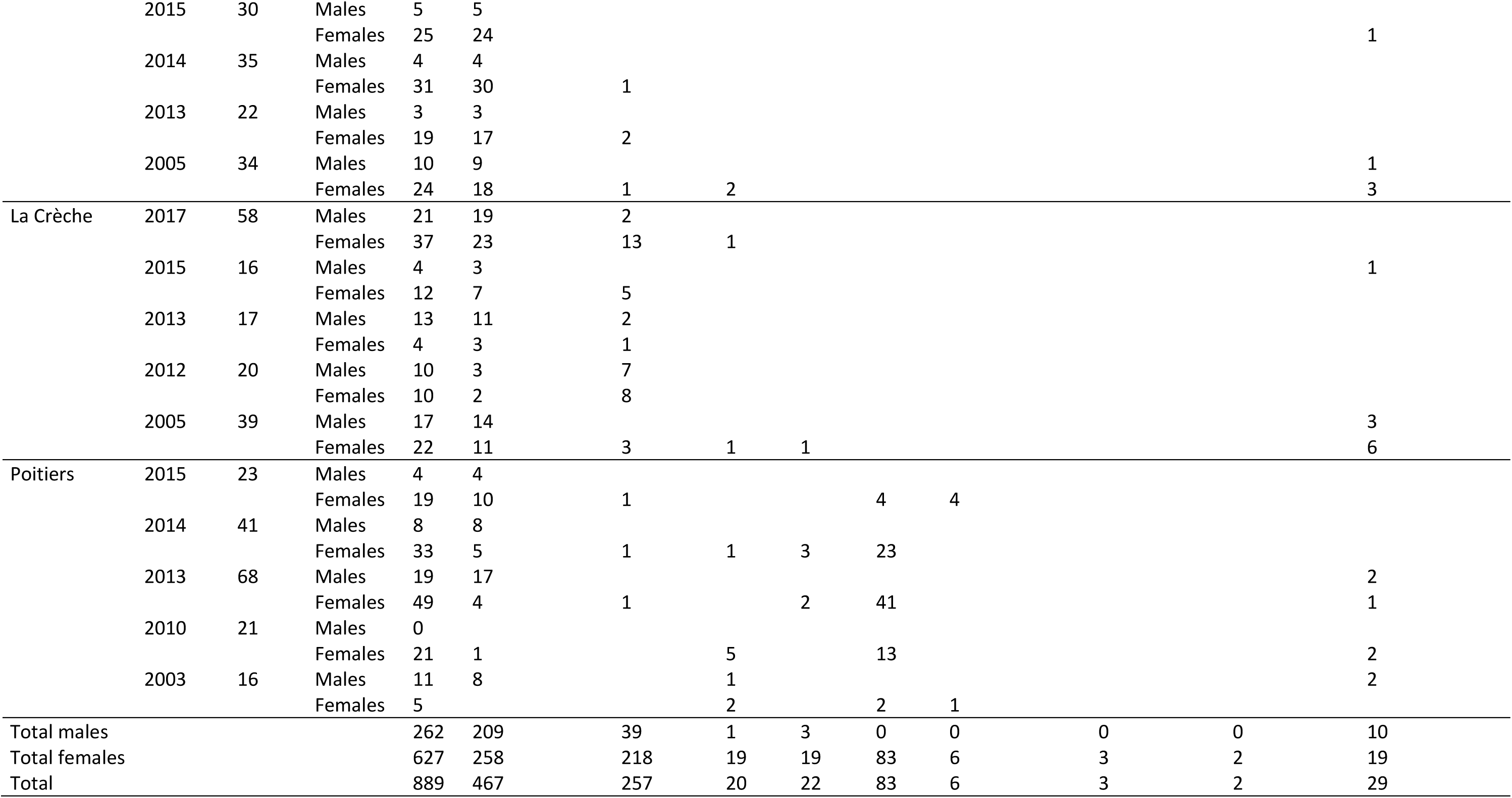
Prevalence of *Wolbachia* and *f* element sex ratio distorters in six populations of *Armadillidium vulgare*.

## Discussion

In this study, we analyzed the temporal dynamics of two SRD segregating in natural populations of the terrestrial isopod *A. vulgare*, to shed light on the potential impact of SRD on the evolution of host sex determination mechanisms at a within-species scale. *Wolbachia* and *f* element distributions were highly heterogeneous among populations in our study area, despite their modest geographic distance of at most 80 km. The emerging trend is that when one SRD is frequent in a population, the other SRD is rare, as noted previously (Durand *et al*., 2023). As it causes a stronger bias toward females, *Wolbachia* is expected to prevail over the *f* element in *A. vulgare* populations (Rigaud, 1997; Cordaux and Gilbert, 2017). Yet, the *f* element was the dominant SRD in four out of six populations we studied and was rising in frequency in Niort (Juchault *et al*., 1992). Overall, the *f* element is more widespread than *Wolbachia* in *A. vulgare* populations (Juchault *et al*., 1993; Durand *et al*., 2023).

Previously proposed explanations include a higher fitness cost entailed by *Wolbachia* relative to the *f* element and occasional paternal transmission of the *f* element (which some males carry, Figure 2) enabled by masculinizing epistatic alleles (Juchault *et al*., 1992; Rigaud and Juchault, 1993; Rigaud, 1997; Rigaud and Moreau, 2004; Cordaux and Gilbert, 2017).

In contrast with spatial heterogeneity, our sampling scheme with up to six sampling time points spanning 12 years per population highlights a global temporal stability in SRD prevalence within populations. At first glance, this qualitative pattern differs from that previously reported for the Niort population, in which *Wolbachia* prevalence was found to decrease concomitantly to an increase of *f* element prevalence over a period of 23 years (Juchault *et al*., 1992). Given *A. vulgare*’s generation time of one year, our study might have spanned too few generations (12) to capture variation in SRD prevalence, which the Niort study spanning 23 generations did. Nevertheless, the time scale of our study enabled us to detect variation in *Wolbachia* strain prevalence, as well as mitochondrial and nuclear variation within and between populations, suggesting that lack of resolution is not an issue. Alternatively, most of the populations we studied may reflect some relatively stable equilibrium with respect to SRD evolutionary dynamics, an equilibrium that the Niort population might not have reached. Indeed, theoretical models have indicated that when feminizing factors are in competition, the one that induces the strongest bias toward females is expected to spread in the population (Taylor, 1990). Thus, a single SRD should remain in the population at equilibrium. Consistently, in most of the populations we analyzed, a single SRD occurs at high frequency, suggesting that these populations may be at or near equilibrium for an SRD. This signal of temporal stability is reminiscent of that recorded in the butterfly *E. mandarina*, in which the feminizing *w*Fem *Wolbachia* strain has been stably maintained at high frequency for 12 years in a Japanese population (Kageyama *et al*., 2020). Similarly, the cytoplasmic incompatibility-inducing *w*Ri *Wolbachia* strain has apparently reached an equilibrium frequency at ∼93% in *D. simulans* populations from California (Weeks *et al*., 2007; Carrington *et al*., 2011). It has been suggested that an apparent stability could be due to hidden processes such as population structure (including extinction-recolonization processes), intragenomic conflicts and coevolutionary processes (Hatcher, 2000).

The temporal stability of SRD in most of *A. vulgare* populations is also reflected in host mitochondrial and nuclear variation, with two notable exceptions. The first one is La Crèche population in 2005, which differs from the other time points (2012 to 2017) on both mitochondrial and nuclear grounds. Interestingly, the sampling spot in La Crèche has been altered by land remodeling between 2005 and 2012. This anthropogenic activity may have caused the reduction or collapse of the historic *A. vulgare* population and the introduction of new individuals as part of the addition of materials during the remodeling (e.g., soil from another location). Such an extinction-recolonization scenario may explain the loss of *Wolbachia* (present at low frequency in 2005) and the increase in *f* element frequency in 2012. It is noteworthy that from 2012 on, SRD, mitochondrial and nuclear variation have been stable, suggesting that stabilization of the population dynamics may be reached in a few years.

The second case of instability is the Poitiers population, which is stable with respect to nuclear variation but not to both mitochondrial and SRD variation. Poitiers is the only population in our dataset in which *Wolbachia* is the dominant SRD across time points, thus highlighting a qualitative pattern of temporal stability. However, our results indicate that the rise in frequency of the *w*VulP strain correlates with a decrease of the *w*VulC strain, suggesting a *Wolbachia* strain replacement in this population. The *w*VulP strain is characterized by a recombination event involving *w*VulC (Verne *et al*., 2007), indicating that *w*VulC is older than *w*VulP, which is consistent with the situation recorded in Poitiers. Assuming the driver of this replacement is *Wolbachia* and not another cytoplasmic element (like the mitochondrion), replacement of *w*VulC by *w*VulP could be due to the latter strain having a transmission advantage over the former strain. Unfortunately, the *w*VulP strain is not very well characterized, and while feminization induction is likely (Verne *et al*., 2007), it has not been formally demonstrated and compared to feminization induced by *w*VulC (Rigaud *et al*., 1991; Cordaux *et al*., 2004). Neither has the respective costs of these two *Wolbachia* strains been investigated. In any event, because *Wolbachia* and mitochondria are co-inherited cytoplasmic entities, changes in *Wolbachia* strains associated with different mitotypes are expected to lead to concomitant changes in mitochondrial variation, but no change in nuclear variation. Therefore, our observations in Poitiers may constitute a typical example of mitochondrial sweep caused by endosymbiont rise in frequency (Galtier *et al*., 2009).

*Wolbachia* dynamics in Poitiers also illustrates that transovarial, maternal transmission is the main transmission mode of *Wolbachia* in *A. vulgare*. However, non-maternal transmission may also occur, as suggested by two individuals with *w*VulM from Beauvoir and La Crèche. These individuals carry mitotypes I and III, respectively, unlike all other *w*VulM-infected individuals which carry mitotype II. As mitotypes I, II and III are distantly related, a plausible explanation is that the two unusual individuals have acquired *Wolbachia* by horizontal transfer, although the hypothesis of historical infections cannot be formally discarded. Horizontal transfer of *Wolbachia* is largely documented in arthropods (O’Neill *et al*., 1992; Werren *et al*., 1995; Heath *et al*., 1999; Vavre *et al*., 1999), including terrestrial isopods (Bouchon *et al*., 1998; Cordaux *et al*., 2001, 2012). Potential mechanisms in isopods include contact between wounded individuals (Rigaud and Juchault, 1995) and cannibalism/predation (Le Clec’h *et al*., 2013). In total, our results suggest that two out of 136 *Wolbachia*-infected individuals could conceivably have acquired their symbionts by horizontal transmission. This may be an underestimate, as horizontal transfers between individuals carrying the same mitotype cannot be detected with our approach. If so, horizontal transmission may occur at a measurable rate in *A. vulgare*, suggesting that it is a parameter of importance in *Wolbachia* evolutionary dynamics in this species.

To conclude, the evolutionary dynamics of SRD, mitochondrial and nuclear variation from various populations over a period of up to 12 years revealed that distributions of *Wolbachia* and the *f* element were much more variable spatially than temporally. This conclusion is well supported for the *f* element, with four populations independently showing the same pattern. For *Wolbachia*, it has remained at high frequency over time but strain genotyping identified an apparently ongoing strain replacement. It is however more difficult to draw strong conclusions for this SRD as it occurred at high frequency in a single population. Such geographic and temporal distribution suggests that migration may not heavily influence SRD evolution in *A. vulgare*, a species exhibiting strong female phylopatry (Durand *et al*., 2019). This is in contrast with the highly dispersive *H. bolina*, in which *Wolbachia* frequency has been shown to fluctuate over time (Hornett *et al*., 2009). Overall, our results provide an empirical basis for future studies on SRD evolutionary dynamics in the context of multiple sex determination factors co-existing within a single species, such as modelling investigations. These efforts will ultimately contribute to assess the impact of SRD on the evolution of host sex determination mechanisms and sex chromosomes.

## Acknowledgments

We thank Sébastien Verne and Victorien Valette for assistance with sample collection. This work was funded by Agence Nationale de la Recherche Grants ANR-15-CE32-0006 (CytoSexDet) to RC and TR, and ANR-20-CE02-0004 (SymChroSex) to JP, and intramural funds from the CNRS and the University of Poitiers.

## Author contribution statement

RC, JP and TR designed the experiments. FG, NB and JP sampled populations. SD, IG and AL performed laboratory work. SD, RP and NB performed data analyses. RC and SD wrote the first draft of the manuscript. TR, JP, NB, RP and FG amended the manuscript.

## Data archiving

Mitotypes are available in GenBank under accession numbers OR074129 to OR074131. All other data are provided in the supplementary information.

