## Supplementary material for "Temporal stability of sex ratio distorter prevalence in natural populations of the isopod *Armadillidium vulgare*": Figs S1-S2 and Tables S1-S5

Content of the file: Figures S1 and S2; Tables S1, S2, S4 and S5.

Table S3 is provided as a separate .xlsx file due to its large size.

**Figure S1.** Variation in mitotype counts across years for three *Armadillidium vulgare* populations from Beauvoir, Chizé and Gript. Prevalence of sex ratio distorters is coded as indicated in the key on the right-hand side of the figure.

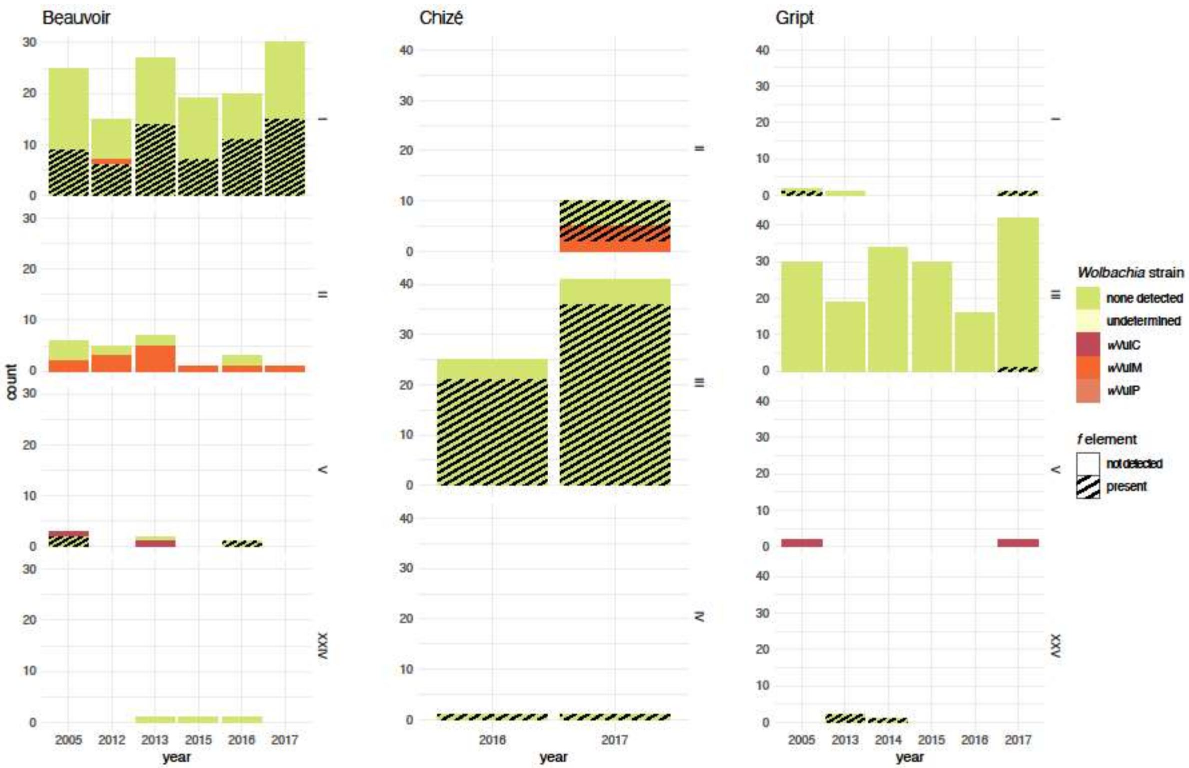

**Figure S2.** Graph of Evanno's delta-K statistic for 2 to 9 simulated genetic clusters (K).

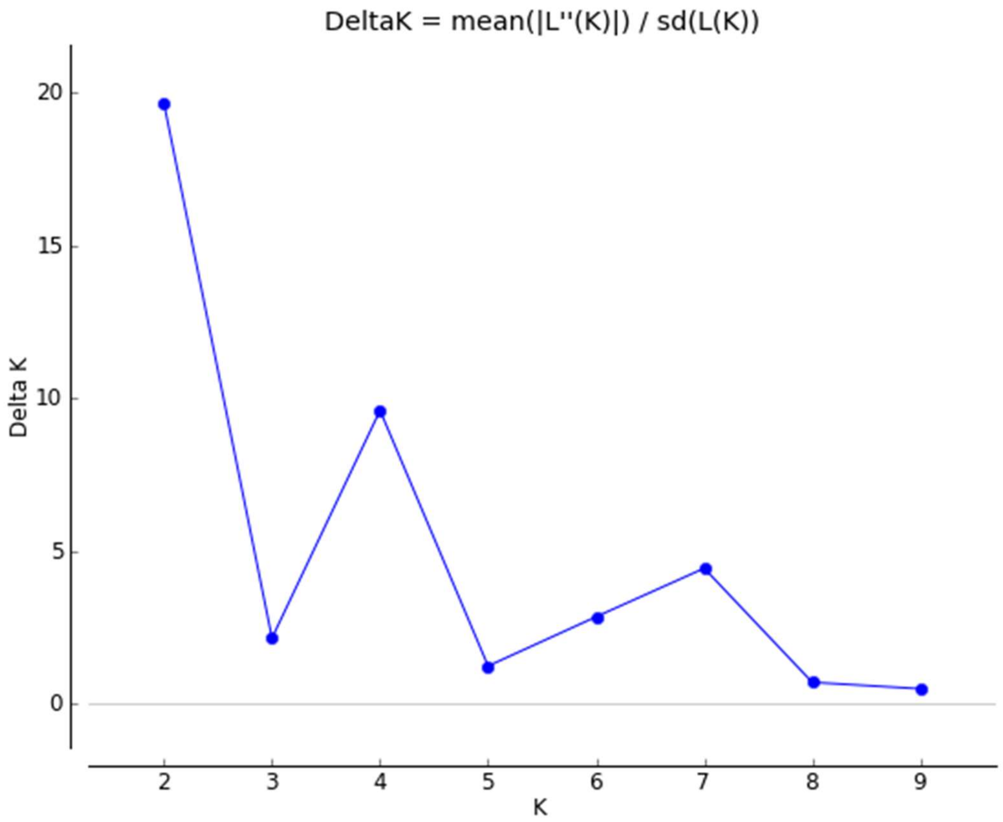

**Table S1.** Molecular markers used for mitochondria, *Wolbachia* and the *f* element.

| Marker name | Target | Primer name | Primer sequence 5'-3' | Tm (°C) | PCR product size (bp) | Reference |
|---|---|---|---|---|---|---|
| *COI* | mitochondria | LCO | GGTCAACAAATCATTAAAGATATTGG | 52 | 700 | Folmer et al. (1994) |
| | | HCO | TAAACTTCAGGGTGACCAAAAAATCA | | | |
| *Jtel* | *f* element | SubF1 | ACGAAAAGCCGACGTAAAATATTT | 55 | 700 | Leclercq et al. (2016) |
| | | JTelR2 | GAAATAAAAGAGCCTGACT | | | |
| *wsp* | *Wolbachia* | 81F | TGGTCCAATAAGTGATGAAGAAAC | 55 | 650 | Braig et al. (1998) |
| | | 691R | AAAAATTAAACGCTACTCCA | | | |
| *recR* | *Wolbachia* | RecR_F_B2 | TGCTTTTCTTCATTTGTTTCCA | 60 | 876 | Badawi et al. (2014) |
| | | RecR_R_B2 | TTTCTAGGCTGAAGTATGCCACT | | | |
| *ftsZ* | *Wolbachia* and *f* element | ftsZf1 | GTTGTCGCAAATACCGATGC | 47 | 1000 | Werren et al. (1995) |
| | | ftsZr1 | CTTAAGTAAGCTGGTATATC | | | |

**Table S2.** Molecular markers used for microsatellite genotyping.

| Multiplex (reference) | Marker name | Primer sequence 5'-3' | Forward primer dye | Repeat type | Observed size range (bp) / scoring issue |
|---|---|---|---|---|---|
| M1 (Verne et al. 2006) | Av1 | TGGAGTCAACTCACATTCTG<br>TGTCTGTAAAACTTGTGCTACG | 6_FAM | (CA)n | 109-116 |
| | Av2 | TGAAGTTCGGGTGAATTGTG<br>ATACCATGACGTGTCGCAAG | 6_FAM | (CA)n(CG)n(CA)n | 154-180 |
| | Av4 | CCGAACCTTTCGAAGGTATT<br>AAGGCACATAACATTTTCACAAA | 6_FAM | (GT)nAT(GT)n(GA)n AA(GA)n | multiple peaks |
| | Av5 | CGTGCGAAGTTCAGATTCTTT<br>GCGCGCTCGAGGATTTAC | 6_FAM | (CA)n | multiple peaks |
| | Av9 | TCTCGAAGAATTGCCTCACA<br>CGATGACTGGGACAATCTCA | HEX | (CA)n | 171-221 |
| M2 (Verne et al. 2006) | Av6 | GGAATGAGGTCGTCGACTATG<br>GTCTTTCAAACGGGCACAAT | 6_FAM | (GT)nAT(GT)n | 167-181 |
| | Av3 | TGAGTCTCATTATAGTTTGGATGA<br>TCCTCTCTATACCCCATAATTTCA | 6_FAM | (CA)n(CG)n(CA)n | 182-236 |
| | Av8 | CAACATCCTAATATTCAGTTCTCA<br>AGTTTACAGAATACGGCTGAGG | HEX | (CA)n | 147-191 |
| M3 (Giraud et al. 2013) | AV0023 | TGGAATTTATGTTTGGAGAGGG<br>GAGGTTAAGTCTGGGGTCGG | HEX | (AG)n | 193 / monomorphic |
| | AV0056 | TTCAAAGGAGCGTTTGACCT<br>AACCACAGCAACAACAGCAG | 6_FAM | (GTT)n | 199-226 |
| | AV0085 | CATGCCGTAAGTCCTCTAGACA<br>TGTGTTATGGTAATTACATTGAAGTTT | NED | (GTA)n | 169-190 |
| | AV0086 | CCCTTGGCTTCCGATACTT<br>TGTCCACAAAGCCAAAATGA | HEX | (TTC)n | 110-122 |
| | AV0096 | TGGCATAAACCAGCTATAAACC<br>TAGTTGCTTTTCCCCTACTTTTG | 6_FAM | (AAC)n | inconsistent amplification |
| M4 (Giraud et al. 2013) | AV0002 | CGACTCCGACTCCGAATG<br>TTCCGACATGTACGATTTTATCA | 6_FAM | (ACTCCG)n | 260-308 |
| | AV0016 | GCTCATTTATGATCTCGTCGC<br>CTCCCACGTGGTTGATCTTC | 6_FAM | (TC)n | 114-126 |
| | AV0018 | GAAGAAATTCAAACTTCACCATCA<br>CTTTGAACAGACTTACGAATAACATC | HEX | (CAA)n | 97-136 |
| | AV0032 | TTTCAACCTTCCTAACCAAACC<br>TTGTTTTATATCCACGACCATCC | NED | (TC)n | 89-105 |
| | AV0099 | CCCCATTTGTGCATGTAGTG<br>ACCCCTCGCTTACATTACCC | HEX | (TG)n | 160-205 / null alleles |
| M5 (Giraud et al. 2013) | AV0061 | GTTTGTATGCATTTACCCCTCTTC<br>GTATGGAACGAAGGGACCG | HEX | (CT)n | 140 / monomorphic |
| | AV0063 | CAAAACATCTGTACGGATTCCC<br>GCCAAACATAAATGCTCGCT | NED | (TACA)n | 126-138 |
| | AV0089 | TTGTTACTTCTACCACCACTATTGC<br>TGGCTCTATAATGATCAATGGAA | HEX | (CTA)n | 86-98 |
| | AV0128 | TGTCGTTGTGAACAGGCTAAA<br>CGTCCGTCGAATGATATTTGT | 6_FAM | (GAT)n | inconsistent amplification |

**Table S4.** Allelic richness (A) and heterozygosity (B) for 28 subpopulations from six French *Armadillidium vulgare* populations and 15 microsatellite loci included in this study.

(A)

| Population | Beauvoir | | | | | | Chizé | | Coulombiers | | | | | Gript | | | | | | La Crèche | | | | | Poitiers | | | | All |
|---|---|---|---|---|---|---|---|---|---|---|---|---|---|---|---|---|---|---|---|---|---|---|---|---|---|---|---|---|---|
| Year | 2005 | 2012 | 2013 | 2015 | 2016 | 2017 | 2016 | 2017 | 2005 | 2014 | 2015 | 2016 | 2017 | 2005 | 2013 | 2014 | 2015 | 2016 | 2017 | 2005 | 2012 | 2013 | 2015 | 2017 | 2003 | 2010 | 2013 | 2014 | population |
| Av1 | 3.19 | 2.84 | 3.36 | 3.21 | 3.22 | 3.22 | 3.31 | 2.94 | 2.19 | 2.66 | 2.45 | 2.53 | 2.43 | 3.25 | 3.43 | 3.25 | 3.25 | 3.21 | 3.24 | 3.13 | 2.93 | 2.96 | 2.93 | 2.77 | 1.85 | 2.46 | 2.40 | 2.17 | 3.10 |
| Av2 | 3.22 | 3.29 | 3.00 | 3.35 | 3.15 | 3.28 | 3.12 | 3.16 | 3.05 | 3.19 | 2.98 | 3.05 | 2.97 | 2.25 | 2.72 | 2.71 | 2.80 | 2.84 | 2.54 | 2.92 | 2.61 | 2.21 | 2.62 | 2.69 | 3.02 | 2.97 | 2.99 | 3.21 | 2.99 |
| Av9 | 3.84 | 4.63 | 4.32 | 4.09 | 4.33 | 3.90 | 3.94 | 3.96 | 4.55 | 4.28 | 4.38 | 4.49 | 4.46 | 4.28 | 3.39 | 3.93 | 4.23 | 3.96 | 3.79 | 4.29 | 3.85 | 4.03 | 4.50 | 4.52 | 4.35 | 4.30 | 4.49 | 4.78 | 4.38 |
| Av3 | 4.27 | 4.10 | 3.90 | 4.29 | 3.54 | 4.43 | 4.69 | 4.38 | 3.99 | 3.04 | 3.56 | 3.58 | 3.93 | 3.53 | 4.07 | 4.18 | 3.15 | 3.77 | 3.57 | 4.33 | 3.39 | 3.43 | 3.29 | 3.78 | 3.02 | 3.53 | 3.66 | 3.72 | 4.10 |
| Av6 | 3.17 | 3.28 | 2.97 | 3.14 | 2.75 | 2.58 | 2.61 | 2.91 | 2.24 | 2.49 | 3.09 | 2.04 | 2.71 | 2.58 | 2.46 | 2.20 | 2.66 | 2.52 | 2.78 | 2.87 | 2.46 | 3.40 | 2.46 | 2.69 | 2.32 | 2.23 | 2.05 | 2.70 | 2.72 |
| Av8 | 4.64 | 3.90 | 3.89 | 4.41 | 4.45 | 4.19 | 4.12 | 4.19 | 4.05 | 4.07 | 4.41 | 4.28 | 4.07 | 3.68 | 3.27 | 3.27 | 3.57 | 3.71 | 3.56 | 4.31 | 3.98 | 3.37 | 3.74 | 4.36 | 4.24 | 4.22 | 4.62 | 4.40 | 4.20 |
| AV0056 | 3.61 | 2.40 | 3.41 | 3.17 | 3.23 | 3.63 | 2.48 | 2.60 | 2.85 | 3.00 | 2.33 | 2.96 | 3.09 | 3.03 | 2.39 | 2.71 | 2.44 | 2.95 | 2.36 | 2.29 | 2.70 | 2.62 | 2.37 | 2.62 | 3.18 | 2.97 | 2.79 | 3.23 | 2.90 |
| AV0085 | 1.16 | 1.84 | 1.88 | 2.00 | 1.81 | 1.61 | 1.78 | 2.22 | 1.00 | 1.25 | 1.13 | 1.27 | 1.36 | 1.36 | 1.83 | 1.60 | 1.79 | 1.35 | 1.29 | 1.53 | 1.29 | 1.23 | 1.19 | 1.45 | 1.23 | 1.23 | 1.28 | 1.00 | 1.52 |
| AV0086 | 1.45 | 1.64 | 1.48 | 1.16 | 1.56 | 1.21 | 1.60 | 1.50 | 1.25 | 1.00 | 1.00 | 1.00 | 1.00 | 1.55 | 1.55 | 1.92 | 1.91 | 2.08 | 1.74 | 2.07 | 1.32 | 1.46 | 1.38 | 1.86 | 1.00 | 1.23 | 1.35 | 1.19 | 1.60 |
| AV0002 | 1.75 | 2.60 | 2.50 | 2.72 | 1.88 | 2.68 | 2.30 | 1.94 | 2.62 | 2.49 | 2.60 | 2.60 | 2.73 | 1.96 | 3.68 | 2.36 | 2.68 | 2.67 | 2.91 | 2.08 | 2.19 | 1.50 | 2.40 | 2.35 | 2.44 | 2.87 | 2.52 | 2.00 | 2.51 |
| AV0016 | 1.43 | 1.15 | 1.00 | 1.70 | 1.95 | 1.72 | 1.24 | 1.12 | 1.65 | 1.71 | 1.78 | 1.59 | 1.87 | 1.11 | 1.63 | 1.65 | 1.59 | 1.67 | 1.67 | 1.10 | 1.67 | 1.46 | 2.02 | 1.51 | 1.23 | 1.00 | 1.20 | 1.48 | 1.52 |
| AV0018 | 3.24 | 3.50 | 3.09 | 3.23 | 3.35 | 3.35 | 2.92 | 3.16 | 3.16 | 2.75 | 2.34 | 3.06 | 2.94 | 3.17 | 3.27 | 2.92 | 3.07 | 3.13 | 2.82 | 3.36 | 3.13 | 3.24 | 2.74 | 3.17 | 3.51 | 2.97 | 3.17 | 3.20 | 3.21 |
| AV0032 | 1.76 | 1.99 | 1.67 | 1.51 | 1.79 | 1.58 | 2.14 | 2.04 | 1.52 | 1.98 | 1.87 | 1.56 | 1.72 | 1.57 | 1.27 | 1.77 | 1.64 | 1.19 | 1.79 | 1.81 | 2.14 | 1.88 | 1.77 | 2.01 | 1.92 | 1.56 | 1.77 | 1.42 | 1.79 |
| Av0063 | 2.35 | 2.18 | 2.44 | 2.28 | 2.08 | 2.17 | 2.35 | 2.49 | 1.83 | 1.86 | 1.69 | 1.89 | 1.90 | 1.85 | 1.75 | 1.80 | 2.03 | 1.38 | 1.81 | 1.94 | 2.19 | 2.33 | 2.68 | 2.21 | 1.83 | 1.56 | 1.93 | 2.05 | 2.09 |
| AV0089 | 1.16 | 1.00 | 1.00 | 1.16 | 1.32 | 1.21 | 1.24 | 1.36 | 1.00 | 1.25 | 1.43 | 1.38 | 1.46 | 1.10 | 1.27 | 1.10 | 1.42 | 1.19 | 1.46 | 1.00 | 1.29 | 1.00 | 1.19 | 1.07 | 1.23 | 1.56 | 1.74 | 1.42 | 1.28 |
| Mean | 2.68 | 2.69 | 2.66 | 2.76 | 2.69 | 2.71 | 2.66 | 2.66 | 2.46 | 2.47 | 2.47 | 2.48 | 2.58 | 2.42 | 2.53 | 2.49 | 2.55 | 2.51 | 2.49 | 2.60 | 2.48 | 2.41 | 2.48 | 2.60 | 2.42 | 2.44 | 2.53 | 2.53 | 2.66 |

(B)

| Population | Beauvoir | | | | | | Chizé | | Coulombiers | | | | | Gript | | | | | | La Crèche | | | | | Poitiers | | | |
|---|---|---|---|---|---|---|---|---|---|---|---|---|---|---|---|---|---|---|---|---|---|---|---|---|---|---|---|---|
| Year | 2005 | 2012 | 2013 | 2015 | 2016 | 2017 | 2016 | 2017 | 2005 | 2014 | 2015 | 2016 | 2017 | 2005 | 2013 | 2014 | 2015 | 2016 | 2017 | 2005 | 2012 | 2013 | 2015 | 2017 | 2003 | 2010 | 2013 | 2014 |
| Av1 | 0.74 | 0.67 | 0.77 | 0.75 | 0.74 | 0.74 | 0.76 | 0.68 | 0.53 | 0.63 | 0.59 | 0.61 | 0.56 | 0.74 | 0.79 | 0.74 | 0.75 | 0.74 | 0.74 | 0.72 | 0.68 | 0.71 | 0.67 | 0.65 | 0.30 | 0.57 | 0.54 | 0.50 |
| Av2 | 0.69 | 0.71 | 0.71 | 0.72 | 0.68 | 0.70 | 0.68 | 0.68 | 0.69 | 0.74 | 0.69 | 0.70 | 0.66 | 0.55 | 0.66 | 0.66 | 0.66 | 0.67 | 0.62 | 0.67 | 0.61 | 0.55 | 0.62 | 0.61 | 0.69 | 0.70 | 0.68 | 0.74 |
| Av9 | 0.80 | 0.89 | 0.86 | 0.83 | 0.86 | 0.82 | 0.82 | 0.82 | 0.89 | 0.86 | 0.87 | 0.88 | 0.88 | 0.86 | 0.70 | 0.82 | 0.84 | 0.83 | 0.80 | 0.86 | 0.81 | 0.84 | 0.89 | 0.88 | 0.87 | 0.85 | 0.87 | 0.91 |
| Av3 | 0.85 | 0.84 | 0.80 | 0.86 | 0.76 | 0.87 | 0.90 | 0.86 | 0.82 | 0.61 | 0.75 | 0.77 | 0.81 | 0.74 | 0.81 | 0.84 | 0.70 | 0.79 | 0.76 | 0.86 | 0.74 | 0.74 | 0.74 | 0.79 | 0.70 | 0.75 | 0.77 | 0.79 |
| Av6 | 0.68 | 0.71 | 0.66 | 0.69 | 0.57 | 0.51 | 0.61 | 0.66 | 0.43 | 0.53 | 0.66 | 0.37 | 0.56 | 0.59 | 0.47 | 0.42 | 0.57 | 0.51 | 0.59 | 0.60 | 0.48 | 0.74 | 0.48 | 0.56 | 0.45 | 0.45 | 0.34 | 0.55 |
| Av8 | 0.89 | 0.81 | 0.80 | 0.85 | 0.87 | 0.82 | 0.83 | 0.82 | 0.83 | 0.79 | 0.86 | 0.84 | 0.82 | 0.77 | 0.65 | 0.69 | 0.75 | 0.78 | 0.73 | 0.85 | 0.81 | 0.72 | 0.78 | 0.86 | 0.86 | 0.87 | 0.89 | 0.86 |
| AV0056 | 0.74 | 0.46 | 0.73 | 0.66 | 0.70 | 0.77 | 0.51 | 0.53 | 0.61 | 0.64 | 0.45 | 0.60 | 0.66 | 0.65 | 0.46 | 0.60 | 0.51 | 0.65 | 0.47 | 0.43 | 0.57 | 0.51 | 0.49 | 0.53 | 0.70 | 0.64 | 0.64 | 0.69 |
| AV0085 | 0.05 | 0.28 | 0.29 | 0.33 | 0.29 | 0.20 | 0.28 | 0.45 | 0.00 | 0.08 | 0.04 | 0.09 | 0.13 | 0.13 | 0.36 | 0.22 | 0.29 | 0.12 | 0.10 | 0.20 | 0.10 | 0.08 | 0.06 | 0.15 | 0.08 | 0.08 | 0.10 | 0.00 |
| AV0086 | 0.15 | 0.23 | 0.18 | 0.05 | 0.22 | 0.07 | 0.20 | 0.18 | 0.09 | 0.00 | 0.00 | 0.00 | 0.00 | 0.20 | 0.18 | 0.33 | 0.36 | 0.46 | 0.30 | 0.53 | 0.10 | 0.15 | 0.12 | 0.31 | 0.00 | 0.08 | 0.13 | 0.07 |
| AV0002 | 0.32 | 0.52 | 0.52 | 0.58 | 0.36 | 0.57 | 0.43 | 0.32 | 0.64 | 0.63 | 0.61 | 0.64 | 0.66 | 0.31 | 0.83 | 0.55 | 0.60 | 0.58 | 0.67 | 0.36 | 0.43 | 0.16 | 0.51 | 0.48 | 0.55 | 0.72 | 0.56 | 0.50 |
| AV0016 | 0.16 | 0.05 | 0.00 | 0.29 | 0.37 | 0.26 | 0.08 | 0.04 | 0.27 | 0.29 | 0.34 | 0.20 | 0.41 | 0.04 | 0.23 | 0.27 | 0.23 | 0.28 | 0.28 | 0.03 | 0.24 | 0.15 | 0.38 | 0.18 | 0.08 | 0.00 | 0.07 | 0.18 |
| AV0018 | 0.73 | 0.77 | 0.73 | 0.73 | 0.75 | 0.75 | 0.68 | 0.72 | 0.70 | 0.61 | 0.45 | 0.68 | 0.66 | 0.72 | 0.73 | 0.66 | 0.69 | 0.69 | 0.66 | 0.75 | 0.72 | 0.75 | 0.62 | 0.71 | 0.78 | 0.67 | 0.71 | 0.73 |
| AV0032 | 0.32 | 0.38 | 0.27 | 0.19 | 0.31 | 0.20 | 0.44 | 0.45 | 0.20 | 0.50 | 0.40 | 0.21 | 0.31 | 0.19 | 0.09 | 0.34 | 0.26 | 0.06 | 0.30 | 0.29 | 0.45 | 0.41 | 0.28 | 0.41 | 0.45 | 0.21 | 0.34 | 0.16 |
| Av0063 | 0.55 | 0.50 | 0.60 | 0.50 | 0.45 | 0.49 | 0.56 | 0.60 | 0.37 | 0.39 | 0.28 | 0.42 | 0.42 | 0.39 | 0.31 | 0.32 | 0.43 | 0.12 | 0.30 | 0.41 | 0.43 | 0.50 | 0.65 | 0.45 | 0.38 | 0.21 | 0.37 | 0.45 |
| AV0089 | 0.05 | 0.00 | 0.00 | 0.05 | 0.12 | 0.07 | 0.08 | 0.12 | 0.00 | 0.08 | 0.16 | 0.14 | 0.17 | 0.03 | 0.09 | 0.03 | 0.16 | 0.06 | 0.16 | 0.00 | 0.10 | 0.00 | 0.06 | 0.02 | 0.08 | 0.22 | 0.32 | 0.16 |
| Mean | 0.52 | 0.52 | 0.53 | 0.54 | 0.54 | 0.52 | 0.52 | 0.53 | 0.47 | 0.49 | 0.48 | 0.48 | 0.51 | 0.46 | 0.49 | 0.50 | 0.52 | 0.49 | 0.50 | 0.50 | 0.48 | 0.47 | 0.49 | 0.51 | 0.46 | 0.47 | 0.49 | 0.48 |

**Table S5.** Pairwise Fst values for each population comparison (below diagonal), and their significance (above diagonal). P-value threshold was adjusted with the Bonferroni correction, P=0.0001. *: significant; NS: non significant.

| Populations | | Beauvoir | | | | | | Chizé | | Coulombiers | | | | | Gript | | | | | | La Crèche | | | | | Poitiers | | | |
|---|---|---|---|---|---|---|---|---|---|---|---|---|---|---|---|---|---|---|---|---|---|---|---|---|---|---|---|---|---|
| | **Year** | **2005** | **2012** | **2013** | **2015** | **2016** | **2017** | **2016** | **2017** | **2005** | **2014** | **2015** | **2016** | **2017** | **2005** | **2013** | **2014** | **2015** | **2016** | **2017** | **2005** | **2012** | **2013** | **2015** | **2017** | **2003** | **2010** | **2013** | **2014** |
| **Beauvoir** | **2005** | | NS | NS | NS | NS | NS | NS | * | * | * | * | * | * | * | NS | NS | NS | NS | * | * | * | NS | * | * | * | NS | * | NS |
| | **2012** | 0.000 | | NS | NS | NS | NS | NS | NS | * | * | * | * | * | * | NS | NS | NS | * | * | * | * | NS | * | * | * | NS | * | NS |
| | **2013** | 0.000 | 0.000 | | NS | NS | NS | NS | NS | * | * | * | * | * | NS | NS | NS | NS | * | * | * | * | NS | NS | * | * | NS | * | NS |
| | **2015** | 0.000 | 0.000 | 0.000 | | NS | NS | NS | NS | * | * | * | * | * | * | NS | NS | NS | * | * | * | NS | NS | NS | * | * | NS | * | NS |
| | **2016** | 0.000 | 0.005 | 0.003 | 0.000 | | NS | * | * | * | * | * | * | * | * | NS | NS | * | * | * | * | * | * | * | * | * | NS | * | NS |
| | **2017** | 0.000 | 0.009 | 0.000 | 0.000 | 0.003 | | * | * | * | NS | * | * | * | * | NS | NS | * | * | * | * | * | NS | * | * | * | NS | * | NS |
| **Chizé** | **2016** | 0.013 | 0.009 | 0.006 | 0.009 | 0.026 | 0.022 | | NS | * | * | * | * | * | * | NS | NS | * | * | * | * | * | NS | NS | * | * | NS | * | NS |
| | **2017** | 0.021 | 0.011 | 0.012 | 0.014 | 0.030 | 0.034 | 0.000 | | * | * | * | * | * | * | NS | * | * | * | * | * | * | * | * | * | * | * | * | * |
| **Coulombiers** | **2005** | 0.063 | 0.060 | 0.061 | 0.043 | 0.061 | 0.051 | 0.065 | 0.079 | | NS | NS | NS | NS | * | NS | * | * | * | * | * | * | * | * | * | NS | NS | * | NS |
| | **2014** | 0.057 | 0.066 | 0.075 | 0.065 | 0.070 | 0.075 | 0.070 | 0.083 | 0.022 | | NS | NS | NS | * | NS | NS | * | NS | * | * | * | NS | * | * | NS | NS | NS | NS |
| | **2015** | 0.062 | 0.058 | 0.072 | 0.052 | 0.057 | 0.064 | 0.068 | 0.081 | 0.015 | 0.008 | | NS | NS | * | NS | * | * | * | * | * | * | * | * | * | * | NS | * | NS |
| | **2016** | 0.041 | 0.039 | 0.044 | 0.031 | 0.031 | 0.036 | 0.053 | 0.064 | 0.011 | 0.019 | 0.010 | | NS | * | NS | NS | NS | * | * | * | * | * | * | * | NS | NS | NS | NS |
| | **2017** | 0.055 | 0.056 | 0.058 | 0.039 | 0.047 | 0.054 | 0.069 | 0.080 | 0.002 | 0.004 | 0.001 | 0.001 | | * | NS | NS | * | * | * | * | * | * | * | * | NS | NS | * | NS |
| **Gript** | **2005** | 0.021 | 0.019 | 0.012 | 0.022 | 0.015 | 0.026 | 0.027 | 0.035 | 0.066 | 0.091 | 0.066 | 0.056 | 0.064 | | NS | NS | NS | NS | * | * | * | NS | * | * | * | NS | * | * |
| | **2013** | 0.037 | 0.030 | 0.042 | 0.032 | 0.030 | 0.038 | 0.039 | 0.056 | 0.051 | 0.049 | 0.074 | 0.037 | 0.047 | 0.043 | | NS | NS | NS | NS | NS | NS | NS | NS | NS | NS | NS | NS | NS |
| | **2014** | 0.024 | 0.022 | 0.031 | 0.021 | 0.023 | 0.029 | 0.030 | 0.049 | 0.025 | 0.024 | 0.031 | 0.027 | 0.021 | 0.018 | 0.004 | | NS | NS | NS | NS | NS | NS | NS | NS | * | NS | NS | NS |
| | **2015** | 0.013 | 0.010 | 0.011 | 0.014 | 0.017 | 0.019 | 0.025 | 0.038 | 0.047 | 0.062 | 0.050 | 0.042 | 0.045 | 0.016 | 0.012 | 0.004 | | NS | NS | * | * | NS | NS | * | * | NS | * | NS |
| | **2016** | 0.038 | 0.036 | 0.040 | 0.031 | 0.024 | 0.038 | 0.051 | 0.068 | 0.033 | 0.054 | 0.039 | 0.035 | 0.030 | 0.016 | 0.015 | 0.000 | 0.006 | | NS | * | * | NS | NS | * | NS | NS | * | NS |
| | **2017** | 0.038 | 0.033 | 0.035 | 0.026 | 0.039 | 0.038 | 0.042 | 0.058 | 0.030 | 0.058 | 0.038 | 0.045 | 0.036 | 0.028 | 0.024 | 0.011 | 0.001 | 0.004 | | * | * | * | * | * | * | NS | * | NS |
| **La Crèche** | **2005** | 0.055 | 0.042 | 0.047 | 0.051 | 0.050 | 0.057 | 0.044 | 0.058 | 0.068 | 0.089 | 0.075 | 0.064 | 0.071 | 0.049 | 0.059 | 0.039 | 0.050 | 0.042 | 0.063 | | * | * | * | * | * | NS | * | NS |
| | **2012** | 0.019 | 0.025 | 0.027 | 0.015 | 0.019 | 0.022 | 0.019 | 0.027 | 0.044 | 0.057 | 0.049 | 0.037 | 0.045 | 0.016 | 0.030 | 0.009 | 0.017 | 0.028 | 0.021 | 0.051 | | NS | NS | NS | * | NS | * | NS |
| | **2013** | 0.024 | 0.020 | 0.020 | 0.024 | 0.022 | 0.032 | 0.016 | 0.024 | 0.068 | 0.084 | 0.054 | 0.062 | 0.062 | 0.009 | 0.056 | 0.027 | 0.020 | 0.031 | 0.028 | 0.043 | 0.000 | | NS | NS | * | NS | * | NS |
| | **2015** | 0.035 | 0.034 | 0.027 | 0.017 | 0.033 | 0.036 | 0.027 | 0.035 | 0.037 | 0.071 | 0.064 | 0.043 | 0.041 | 0.035 | 0.035 | 0.024 | 0.024 | 0.037 | 0.026 | 0.053 | 0.007 | 0.010 | | NS | NS | NS | * | NS |
| | **2017** | 0.025 | 0.025 | 0.033 | 0.019 | 0.025 | 0.031 | 0.020 | 0.033 | 0.044 | 0.058 | 0.050 | 0.034 | 0.046 | 0.026 | 0.029 | 0.013 | 0.022 | 0.026 | 0.027 | 0.038 | 0.000 | 0.002 | 0.005 | | * | NS | * | NS |
| **Poitiers** | **2003** | 0.056 | 0.060 | 0.049 | 0.044 | 0.053 | 0.059 | 0.046 | 0.055 | 0.024 | 0.021 | 0.040 | 0.032 | 0.023 | 0.052 | 0.053 | 0.030 | 0.058 | 0.041 | 0.044 | 0.062 | 0.024 | 0.037 | 0.031 | 0.027 | | NS | NS | NS |
| | **2010** | 0.053 | 0.047 | 0.039 | 0.024 | 0.043 | 0.046 | 0.043 | 0.057 | 0.019 | 0.039 | 0.047 | 0.018 | 0.024 | 0.049 | 0.016 | 0.022 | 0.037 | 0.021 | 0.031 | 0.058 | 0.023 | 0.052 | 0.027 | 0.027 | 0.000 | | NS | NS |
| | **2013** | 0.058 | 0.058 | 0.054 | 0.048 | 0.051 | 0.058 | 0.054 | 0.060 | 0.034 | 0.044 | 0.055 | 0.036 | 0.032 | 0.056 | 0.045 | 0.035 | 0.052 | 0.039 | 0.048 | 0.062 | 0.035 | 0.051 | 0.025 | 0.034 | 0.000 | 0.000 | | NS |
| | **2014** | 0.031 | 0.038 | 0.032 | 0.021 | 0.038 | 0.030 | 0.033 | 0.042 | 0.009 | 0.034 | 0.033 | 0.019 | 0.018 | 0.038 | 0.039 | 0.020 | 0.032 | 0.025 | 0.029 | 0.052 | 0.013 | 0.033 | 0.010 | 0.018 | 0.000 | 0.000 | 0.000 | |
